# Impaired astrocytic Ca^2+^ signalling in awake Alzheimer’s disease transgenic mice

**DOI:** 10.1101/2021.11.21.469461

**Authors:** Knut Sindre Åbjørsbråten, Gry H. E. Syverstad Skaaraas, Céline Cunen, Daniel M. Bjørnstad, Kristin M. Gullestad Binder, Vidar Jensen, Lars N.G. Nilsson, Shreyas B. Rao, Wannan Tang, Gudmund Horn Hermansen, Erlend A. Nagelhus, Ole Petter Ottersen, Reidun Torp, Rune Enger

## Abstract

Increased astrocytic Ca^2+^ signaling related to amyloid plaques has been shown in Alzheimer’s disease mouse models, but to date no reports have characterized behaviorally induced astrocytic Ca^2+^ signalling in such mice without the confounding effects of anesthesia. Here, we employ an event-based algorithm to assess astrocytic Ca^2+^ signals in the neocortex of awake-behaving tg-ArcSwe mice and non-transgenic wildtype littermates while monitoring pupil responses and behavior. We demonstrate an attenuated astrocytic Ca^2+^ response to locomotion and an uncoupling of pupil responses and astrocytic Ca^2+^ signalling in 15-months old plaque-bearing mice. This points to a potential decoupling of neuromodulatory activation and astrocytic Ca^2+^ activity, which may account for some of the cognitive dysfunctions observed in Alzheimer’s disease.

## Introduction

Since astrocytic Ca^2+^ signals were first discovered in the early 1990’s they have been the object of numerous studies exploring their roles in brain physiology and pathophysiology. Importantly, such signals have been shown to occur in response to a wide array of neurotransmitters and to trigger the release of substances that affect neuronal signalling and the vasculature. A growing body of evidence suggests that astrocytic Ca^2+^ signals play important roles in higher brain functions such as memory formation and cortical processing, mediated in part through the neuromodulatory systems of the brain (Adamsky et al. 2018; Kol et al. 2020; Poskanzer and Yuste 2016, 2011; Ye et al. 2020; Paukert et al. 2014).

Astrocytic Ca^2+^ signalling in an Alzheimer’s disease (AD) mouse model was first described by Kuchibhotla et al. 2009 (Kuchibhotla et al. 2009), who found pathological Ca^2+^ waves originating at amyloid plaques, and a general increase in astrocytic Ca^2+^ signalling. Later, Delekate et al. showed plaque-associated astrocytic hyperactivity mediated through activation of metabotropic purine receptors (Delekate et al. 2014). These studies were performed under anesthesia, which severely attenuates physiological Ca^2+^ signals (Thrane et al. 2012).

The field of astrocytic Ca^2+^ signalling is undergoing a revolution as developments in optical imaging and genetically encoded fluorescent sensors now allow us to monitor these signals in awake-behaving mice, without the confounding effects of anesthesia (Srinivasan et al. 2015; Bojarskaite et al. 2020). Such studies have revealed exceedingly rich and complex astrocytic Ca^2+^ signalling ranging from large activations of nearly all astrocytes in a field-of-view (FOV) under locomotion and startle responses due to noradrenergic activity (Ding et al. 2013; Paukert et al. 2014), to small, localized signals occurring spontaneously or as a response to local neuronal activity (Bindocci et al. 2017; Stobart et al. 2018; Srinivasan et al. 2015). New analytical tools now also enable us to accurately quantify and describe these signals (Y. Wang et al. 2019; Bjørnstad et al. 2021).

Here we show by two-photon microscopy that ∼15 month old unanesthetized awake-behaving tg-ArcSwe mice display attenuated behaviorally induced Ca^2+^ signalling in cortical astrocytes during locomotion. These mice carry two mutations in the amyloid precursor protein gene, the Arctic (E693G) and Swedish (KM670/6701NL) mutations, and exhibit amyloid-β deposits, a hallmark of AD (Lord et al. 2006; Yang et al. 2011; Lillehaug et al. 2014; Philipson et al. 2009). As noradrenergic signalling is known to be the most potent trigger of astrocytic Ca^2+^ signalling in startle responses and locomotion, and pupil responses are regarded a faithful, although indirect, readout of noradrenergic signalling in the brain during physiological conditions (Reimer et al. 2016; Zuend et al. 2020; Costa and Rudebeck 2016), we compared the pupil responses to the astrocytic Ca^2+^ signals and found a strong positive correlation in wild-type mice. No such correlation was present in the AD mice. This points to a potential decoupling of neuromodulatory activation and astrocytic Ca^2+^ activity. Such perturbed behaviorally induced astrocytic Ca^2+^ signalling may account for some of the cognitive deficiencies observed in AD patients.

## RESULTS

### Two-photon imaging of awake-behaving tg-ArcSwe mice

To characterize astrocytic Ca^2+^ signalling in awake tg-ArcSwe mice and nontransgenic littermates we employed two-photon microscopy of cortical layer 1–3 astrocytes in the somatosensory cortex expressing GCaMP6f. The glial fibrillary acidic protein (GFAP) promoter was used to target astrocytes (Fig 1A). Amyloid plaques were visualized *in vivo* by methoxy-X04 delivered by intraperitoneal injection (Fig. 1A). Methoxy-X04 enters the brain and specifically stains parenchymal Aβ plaques and cerebrovascular deposits (Klunk et al. 2002), and has been used for *in vivo* imaging in transgenic mice with amyloid plaques (Delekate et al. 2014; Kuchibhotla et al. 2009; Meyer-Luehmann et al. 2008). Imaging was performed at ∼30 Hz frame rate to capture fast populations of astrocytic Ca^2+^ transients with simultaneous surveillance video recording of the mouse behavior, movement of the treadmill as well as pupil dilations and constrictions to monitor the level of arousal (Reimer et al. 2016). The mice were allowed to spontaneously move on a custom-built disc shaped treadmill, and all mice exhibited both periods of quiet wakefulness (absence of locomotion), and running (Supplementary Fig. 1). Astrocytic Ca^2+^ signals were analyzed using a newly developed event-based Ca^2+^ signal analysis toolkit, outlining so-called regions-of-activity (ROAs), combined with manually segmented regions-of-interests (ROIs) outlining astrocytic subcompartments (Bjørnstad et al. 2021; Bojarskaite et al. 2020).

**Figure 1.**
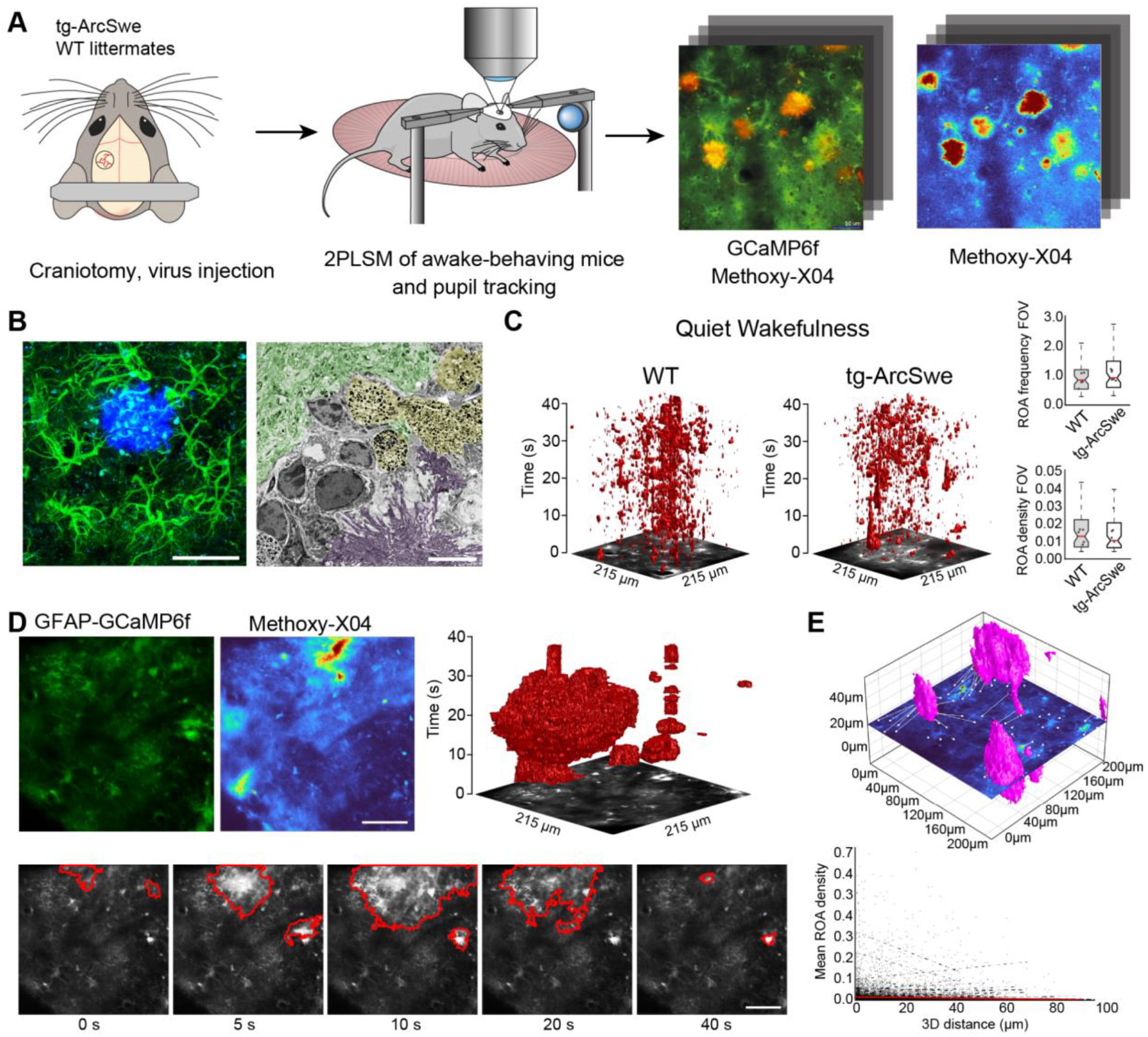
Experimental setup and astrocytic Ca^2+^ signalling in behavioral quiessence. (A) Virus encoding GCaMP6f was injected into the somatosensory cortex of tg-ArcSwe mice and non-mutant littermates. After 3 weeks of recovery, mice were habituated to head-fixation on a disc-shaped treadmill, allowing the mice to move freely at will. Methoxy-X04 was injected 24 hours prior to imaging to visualize amyloid plaques. During imaging, both locomotor activity and pupil responses were recorded. (B) The mice were ∼15 months of age during experiments, at a time when they exhibited dense-core amyloid-β plaques. Left image: confocal micrograph of an amyloid plaque (methoxy-X04, blue) and astrocytes (anti-GFAP, green). Scale bar: 40 μm. Right image: Electron micrograph showing a dense amyloid plaque (purple overlay), autophagic vacuoles (yellow overlay) and relatively normal neuropil morphology (green overlay). Scale bar 2 μm. (C) Astrocytic Ca^2+^ signals during quiet wakefulness (absence of locomotion) in the form of regions-of-activity (ROAs) displayed in an *x-y-t* 3D rendering where red regions denote signal. Box-and-whisker plots representing overall Ca^2+^ signals in quiet wakefulness in tg-ArcSwe mice and littermates. (D) Example of pathological astrocytic Ca^2+^ wave emanating from an amyloid plaque. Top left: average image projections of GCaMP6f fluorescence and methoxy-X04, respectively. Top right: 3D *x-y-t* rendering of ROAs representing a pathological Ca^2+^ wave. Bottom row: micrographs of the same pathological Ca^2+^ wave as in the 3D plot with the Ca^2+^ event outlined in red. Scale bars: 50 μm. (E) 3D visualization of the imaging plane relative to amyloid plaques, with lines representing shortest distance from plaque to ROI. We found a low correlation between distance to nearest plaque and gross level of astrocytic Ca^2+^ signalling.

### Astrocytes close to amyloid plaques express GCaMP6f

The tg-ArcSwe mice were imaged at ∼15 months of age. At this age they present with amyloid-β plaques throughout the cortical mantle, and score poorly on behavioral tasks (Codita et al. 2010; Lillehaug et al. 2014; Lord et al. 2006)(Fig. 1B). Aβ-plaques were characterized by loss of cells and severely perturbed tissue morphology, including autophagic vacuoles (Fig 1B). Even so, relatively normal cellular morphology was present at short distances away from amyloid plaques, and astrocytes faithfully expressed the GCaMP6f Ca^2+^ sensor 3 weeks after viral transduction (Fig. 1A,B).

### Astrocytic Ca^2+^ signals in quiet wakefulness are preserved in tg-ArcSwe mice

In quiet wakefulness (defined as absence of locomotion), we found examples of long-lasting pathological Ca^2+^ waves as reported previously in anesthetized mice (Delekate et al. 2014; Kuchibhotla et al. 2009), often emanating in the vicinity of discernable amyloid plaques and spreading to nearby astrocytes. Such Ca^2+^ waves were found in 10–15% of recordings from tg-ArcSwe mice (Fig. 1D). The number of such clear pathological events were few compared to the overall astrocytic Ca^2+^ signalling we found without the highly confounding effects of anesthesia. Consequently, the gross level of astrocytic Ca^2+^ signalling was similar in mutant mice and their littermates as measured by ROA frequency, ROA density (the active fraction of a compartment) as well as event size and duration in the full field-of-view (FOV) and across the different astrocytic subcompartments (Fig 1C and Supplementary Fig. 2). We were not able to detect a clear correlation in astrocytic Ca^2+^ signalling measured by ROA density and the distance from nearest amyloid plaque (in 3D) (slope = -0.00020, Fig. 1E).

### Uncoupling between pupil dilation and astrocytic Ca^2+^ responses during spontaneous running in tg-ArcSwe mice

Locomotor behavior is known to be strongly correlated with astrocytic Ca^2+^ signalling (Paukert et al. 2014; Bojarskaite et al. 2020; Srinivasan et al. 2015), putatively through the activation of the noradrenergic and cholinergic neuromodulatory systems in conjunction with local network activity (Kjaerby et al. 2017). To investigate if the physiological astrocytic Ca^2+^ responses were preserved in the tg-ArcSwe mice, they were allowed to move freely on a custom built disc-shaped treadmill (Bojarskaite et al. 2020). All mice exhibited both running and behavioral quiessence, and the level of running between the two genotypes were comparable (Supplementary Fig. 1). Running was accompanied by an increase in pupil size and a brisk increase in astrocytic Ca^2+^ signalling typically involving most of the astrocytes in the field-of-view (FOV) in both genotypes (Fig. 2A). When astrocytic Ca^2+^ signals were analysed using a linear mixed effects regression model, a lower ROA density rise rate was found in tg-ArcSwe mice when assessing the full FOV (0.31 in WT vs. 0.20 in tg-ArcSwe, p = 0.032), and astrocytic processes (0.32 in WT vs. 0.20 in tg-ArcSwe, p = 0.032), whereas the Ca^2+^ responses were not significantly different in astrocytic somata and endfeet (0.46 in WT vs. 0.39 in tg-ArcSwe, p = 0.23, and 0.35 in WT vs. 0.30 in tg-ArcSwe, p = 0.11, respectively)(Fig. 2B). Max ROA density values in WT vs. tg-ArcSwe were significantly different when assessing the full FOV (0.63 vs. 0.42, p = 0.033), near significantly different when assessing astrocytic processes and endfeet (0.72 vs. 0.53, p = 0.053 for processes and 0.69 vs. 0.51, p = 0.068 for endfeet), and not significantly different for astrocytic somata (0.81 vs. 0.71, p = 0.25)(Fig. 2B). For mean ROA density values, see Supplementary Fig. 3.

**Figure 2.**
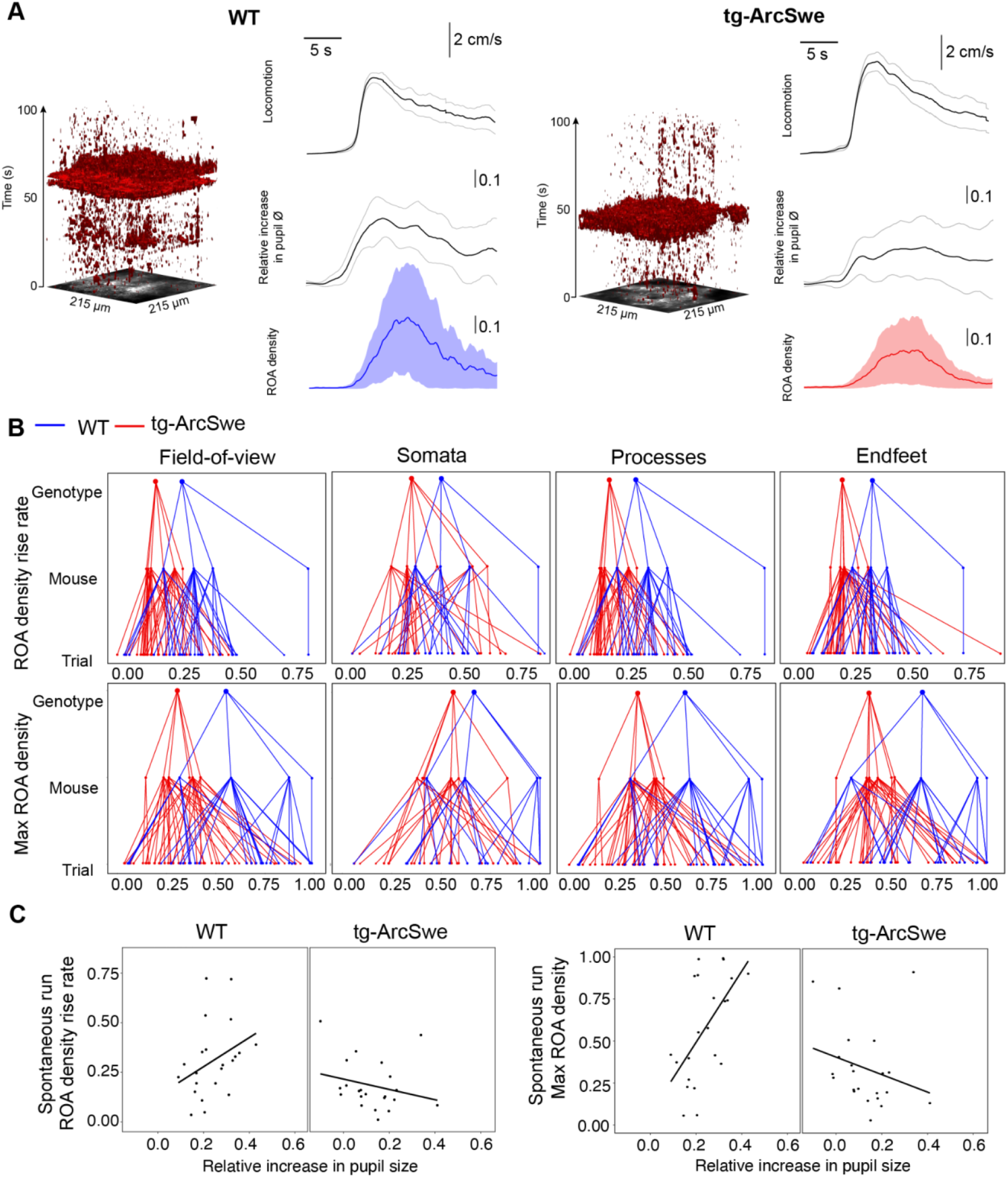
Uncoupling of pupil dilation and astrocytic Ca^2+^ responses during spontaneous running. (A) Bouts of spontaneous running caused a pronounced increase in pupil size and an increase in astrocytic Ca^2+^ signalling. ROAs presented as an *x-y-t* 3D rendering where red regions denote signal. Locomotion trace show mean response across trials with 95% confidence interval, pupil trace and ROA density traces show median across trials ± median absolute deviation. (B) Upper row: Plots showing the median level of ROA density rise rate per genotype (red = tg-ArcSwe, blue = WT littermates), per mouse and per trial in the full FOV, astrocytic somata, astrocytic processes and astrocytic endfeet. Lower row: Same as upper row, but showing median levels of max ROA density in the astrocytic subcompartments. (C) Scatterplots of ROA density rise rate and max ROA density vs. relative increase in pupil size upon spontaneous running.

Pupil responses are known to be a faithful indirect indicator of activity in the locus coeruleus in mice, even though also the cholinergic neuromodulatory system plays a role for sustained pupil dilation (Reimer et al. 2016). As norepinephrine is known to be a potent trigger of astrocytic Ca^2+^ signalling (Bekar, He, and Nedergaard 2008; Srinivasan et al. 2015; Paukert et al. 2014), one would expect to find a strong correlation between pupil dilations and astrocytic Ca^2+^ signals. This was indeed the case for WT mice: When comparing ROA density rise rate and pupil dilation, we found a clear positive slope in line with previous reports (Zuend et al. 2020). In transgenic mice, this correlation was lost, or even reversed (Fig. C, slope of 0.80 in WT vs. -0.35 in tg-ArcSwe, p = 0.007), demonstrating an uncoupling between pupil responses and astrocytic Ca^2+^ signalling in AD transgenic mice. Similarly, when assessing max ROA density, we found a clear positive slope in WT, which was lost in tg-ArcSwe (Fig. 2C, 2.06 in WT vs. -0.58 in tg-ArcSwe, p = 0.00039). Similar slopes were found when comparing mean ROA density vs pupil dilation (Supplementary Fig. 3).

### Uncoupling between pupil dilation and astrocytic Ca^2+^ responses during startle in tg-ArcSwe mice

Another main trigger for astrocytic Ca^2+^ signals are startle responses that are mediated through an activation of the noradrenergic system (Ding et al. 2013; Srinivasan et al. 2015). Even though typically triggering running, the startle response could also trigger freezing behavior and is thought to activate different subcortical networks than spontaneous locomotor behavior (Caggiano et al. 2018; Ferreira-Pinto et al. 2018; Grillner and El Manira 2020). To characterize startle mediated astrocytic activation in the two groups of mice, mice were subjected to 10 air puffs delivered at 10 Hz directed to the vibrissa, nasal and facial region contralaterally to the recording side once per trial at 150 s in a 300 s two-photon imaging recording. Trials in which the mouse was spontaneously running at or immediately before the air puff were excluded from the analyses. We found no signs of habituation to the stimulus in terms of behavioral response (Supplementary Fig. 4). Interestingly, tg-ArcSwe mice were more prone to react with running behavior during startle responses than WT littermates (Fig. 3A), consistent with previous reports of enhanced startle response in other mouse models of AD (McCool et al. 2003). The level of pupil dilation was however similar in the two genotypes (0.17 vs. 0.12 relative increase in pupil size in WT vs. tg-ArcSwe, respectively, p = 0.36, 86 trials)(Fig. 3A). When modelled with a mixed effects linear regression model, for ROA density rise rate, there was a trend of lower Ca^2+^ responses in the full FOV, astrocytic processes and endfeet in tg-ArcSwe mice (0.34 in WT vs. 0.23 in tg-ArcSwe, p = 0.09, 0.42 in WT vs. 0.31, p = 0.09 and 0.36 in WT vs. 0.25, p = 0.10, respectively). No trend was evident in astrocytic somata (0.25 in WT vs. 0.23 in tg-ArcSwe, p = 0.8)(Fig. 3B). Max ROA density was similar for the full FOV (0.36 in WT vs. 0.35 in tg-ArcSwe, p = 0.40), astrocytic somata (0.37 in WT vs. 0.35 in tg-ArcSwe, p = 0.83), astrocytic processes (0.41 in WT vs. 0.34 in tg-ArcSwe, p = 0.39), and astrocytic endfeet (0.41 in WT vs. 0.35 in tg-ArcSwe, p = 0.48). For mean ROA density values, see Supplementary Fig. 3.

**Figure 3.**
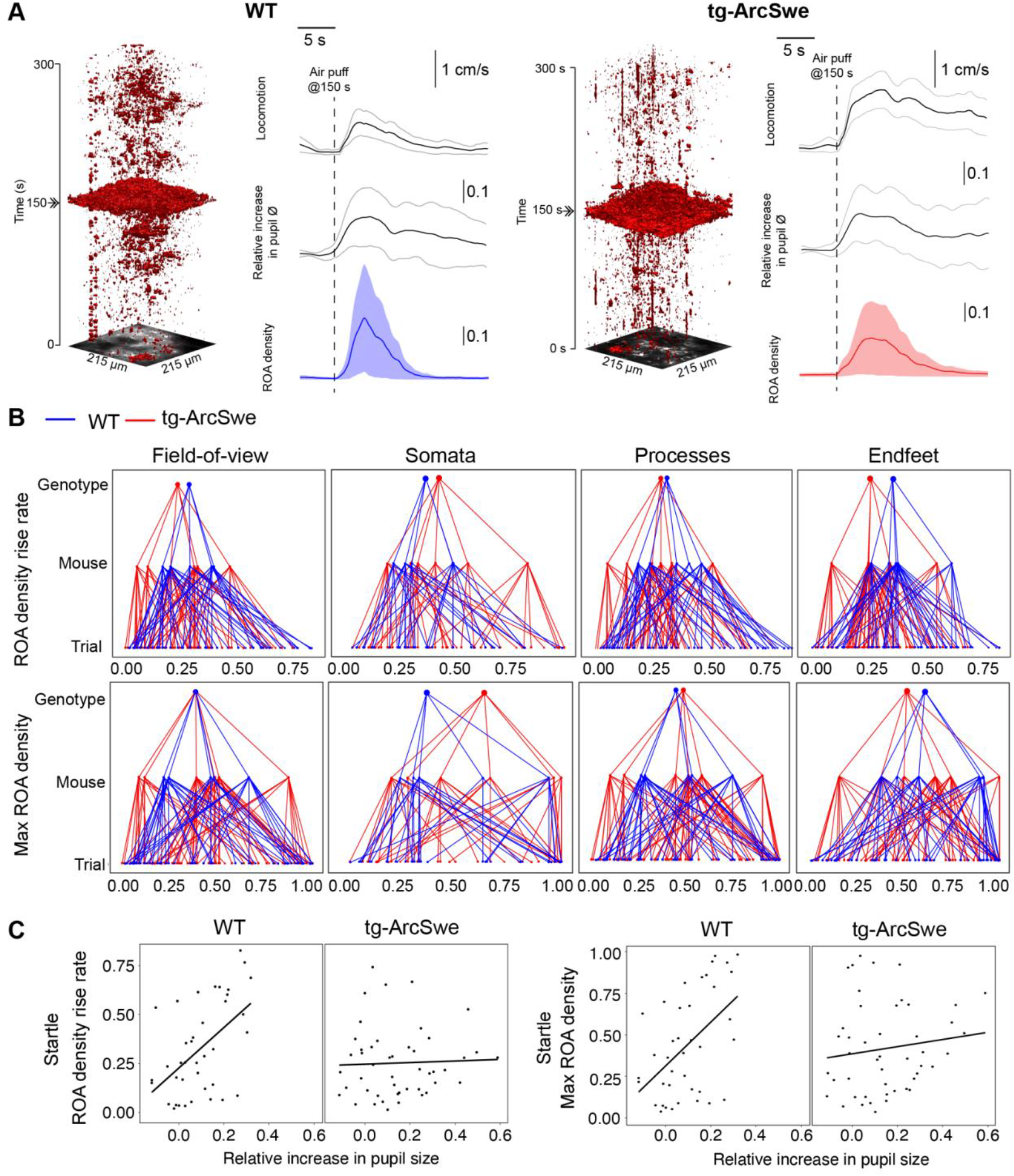
Uncoupling of pupil dilation and astrocytic Ca^2+^ responses during startle. (A) In an imaging trial of 300 s duration, mice were subjected to an air puff to the face/vibrissa contralateral to the imaging window at 150 s. This caused an increase in locomotor activity, pupil dilation, and a pronounced increase in astrocytic Ca^2+^ signalling in both genotypes (right). Locomotion trace show mean response across trials with 95% confidence interval, pupil trace and ROA density traces show median across trials ± median absolute deviation. Astrocytic Ca^2+^ signals in the form of ROAs displayed in an *x-y-t* 3D rendering where red denote signal (left). (B) Upper row: Plots showing the median level of the ROA density rise rate per genotype (red = tg-ArcSwe, blue = WT littermates), per mouse and per trial in the full FOV, astrocytic somata, astrocytic processes and astrocytic endfeet. Lower row: Same as upper row but for max ROA density. (C) Scatterplots of ROA density rise rate and max ROA density vs. relative increase in pupil size.

However, the relationship between astrocytic Ca^2+^ responses and pupillary responses were highly different in the two genotypes (Fig. 3D), with WT mice displaying a clear positive slope, while tg-ArcSwe exhibited a slope close to zero (0.91 vs. -0.013 p = 0.00043 for ROA density rise rate, and 1.27 vs. 0.083 p = 0.0043 and max ROA density, respectively), suggesting an uncoupling between pupillary responses and astrocytic Ca^2+^ responses in tg-ArcSwe mice similar to during spontaneous running (Fig. 3C). Similar slopes were found when comparing mean ROA density vs pupil dilation (Supplementary Fig. 3).

### Widespread reactive astrogliosis in tg-ArcSwe mice

AD transgenic mouse models with high levels of beta-amyloid deposition exhibit reactive astrogliosis (Rodríguez-Arellano et al. 2016), which might affect astrocytic Ca^2+^ activity (Shigetomi et al. 2019; Sano et al., n.d.). We therefore assessed the level of reactive astrogliosis by assessing GFAP expression by immunofluorescence, mRNA levels and morphometry of astrocytes in tg-ArcSwe mice and WT littermates (Fig. 4).

**Figure 4.**
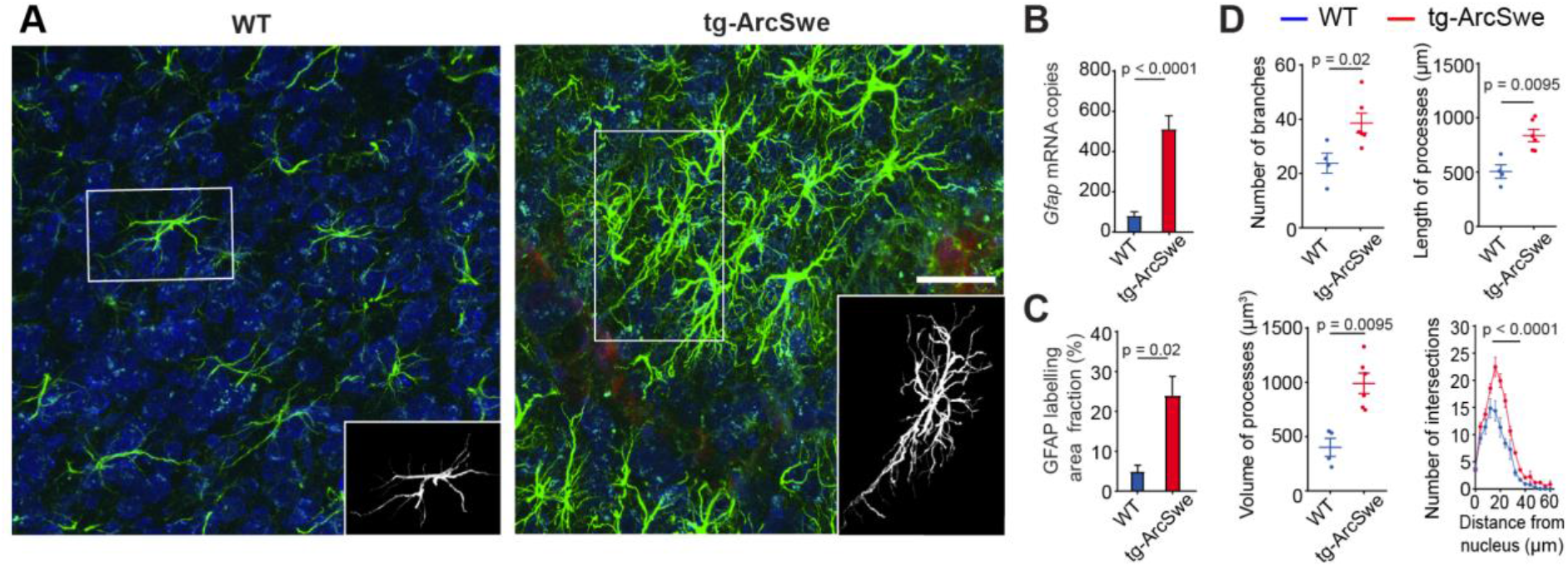
Widespread reactive astrogliosis in tg-ArcSwe mice. (A) Representative micrographs labelled with anti-GFAP antibodies (green) and anti-NeuN antibodies (blue). Scale bar: 40 μm. (B) *Gfap* mRNA expression was considerably higher in tg-ArcSwe mice (p < 0.0001, n = 8 animals in both groups). (C) The area fraction of GFAP labelling was significantly higher in tg-ArcSwe mice compared to controls (n = 6 tg-ArcSwe and 4 WT littermates). (D) Astrocytes were isolated (inset in A) and analyzed with Simple Neurite Tracer (SNT) plugin in FIJI ImageJ. Tg-ArcSwe mice displayed an increase in the total number of branches (p = 0.02), the length and volume of processes (p = 0.0095 and p = 0.095, respectively) of the GFAP labelled astrocytes (n= 8 astrocytes from each animal. 6 tg-ArcSwe and 4 WT littermates), as well as an increased number of branching points and intersections at 16–32 um distance from the nucleus (p < 0.0001).

We found a strong increase in levels of mRNA encoding GFAP (81.58 ± 19.56 for WT; 512.13 ± 66.95 for tg-ArcSwe, p < 0.0001, n = 8 animals in both groups) in tg-ArcSwe compared to WT littermates (Fig. 4C). This was supported by a significantly higher GFAP labeling fraction in tg-ArcSwe animals compared to WT littermates (Fig. 4B, 24.03% ± 4.80% in tg-ArcSwe, n = 6 mice vs. 4.77% ± 1.58% in WT, n = 4 mice, p = 0.02). Morphometric analyses displayed a significantly higher number of labelled astrocytic processes (38.63 ± 3.61 in tg-ArcSwe, n = 6 mice; 23.84 ± 3.72 in WT, n = 4 mice; p = 0.02), as well as total length (837.9 µm ± 55.9 µm for tg-ArcSwe; 506.0 µm ± 62.5 µm for WT; p = 0.0095) and volume of processes (990.9 µm^3^ ± 93.8 µm^3^ for tg-ArcSwe; 401.5 µm^3^ ± 82.9 µm^3^ for WT; p = 0.0095) in tg-ArcSwe mice compared to WT littermates (Fig. 4D). In addition, astrocytes in the tg-ArcSwe mice exhibited significantly more branching points at the distance 16–32 µm from the nucleus (p < 0.0001).

## Discussion

Astrocytic Ca^2+^ signalling is emerging as a key component of signal processing in the brain and figures prominently in brain state transitions and memory formation (Poskanzer and Yuste 2016, 2011; Adamsky et al. 2018; Bojarskaite et al. 2020; Vaidyanathan et al. 2021). Aberrant astrocytic Ca^2+^ signalling could hence be implicated in the perturbed cognition seen in dementia. Indeed, previous studies have shown increased astrocytic Ca^2+^ signalling and spreading pathological Ca^2+^ waves in AD mouse models (Kuchibhotla et al. 2009; Delekate et al. 2014). Based on these studies in anesthetized animals and other studies (Haughey and Mattson 2003; Lim et al. 2013; Abramov, Canevari, and Duchen 2004, 2003; Verkhratsky 2019) the current concept is that aberrant signalling to some degree is spatially coupled to amyloid deposits – the pathological hallmark of Alzheimer’s disease.

Methodological advances now allow astrocytic Ca^2+^ signals to be studied in awake animals. Benefitting from this opportunity we show that tg-ArcSwe mice sustain a pattern of behaviorally induced astrocytic Ca^2+^ signalling similar to that found in littermate controls. However, the signals are weaker than in controls and do not display the correlation with pupil responses typically seen in wild type animals. The behaviorally induced Ca^2+^ signals bear no clear spatial correlation to amyloid plaques. We conclude that elimination of anaesthesia unveils a new dimension of Ca^2+^ signalling, superimposed on the locally induced signals described in previous studies. The uniform attenuation of the behaviorally induced signals in tg-Arc-Swe mice and their uncoupling from pupil responses point to a potential perturbation of neuromodulatory control.

Astrocytic Ca^2+^ signalling in awake behaving mice is dominated by norepinephrine-induced astrocytic Ca^2+^ signals across the cortical mantle in relation to locomotor or startle responses (Srinivasan et al. 2015; Ding et al. 2013). The downstream effects of these synchronized, global Ca^2+^ signals coupled to arousal are not fully understood, but they may play a role in altering the levels of extracellular K^+^ (F. Wang et al. 2012), release of gliotransmitters such as glutamate or ATP (Kjaerby et al. 2017; Haydon and Nedergaard 2014) or metabolic supply (Zuend et al. 2020). Independent of what are the exact downstream mechanisms, there are reasons to believe that astrocytes serve as actuators of the noradrenergic neuromodulatory system, presumably exerting noradrenergic effects on neural network processing and ultimately affecting cognitive function (Ye et al. 2020; Holland, Robbins, and Rowe 2021; Poskanzer and Yuste 2016).

The cause of perturbed astrocytic Ca^2+^ responses during locomotion responses in AD mice in our experiments is not entirely clear. Firstly, we find prominent reactive astrogliosis throughout the brains of the mice we studied. Although still sparsely investigated, both attenuated and increased Ca^2+^ signalling have been demonstrated in various models of reactive astrogliosis, but so far without a clear understanding of the mechanisms involved (Shigetomi et al. 2019). Widespread reactive astrogliosis is nonetheless highly likely to perturb the physiological signalling of astrocytes, as prominent changes in the expression of key receptors and molecules of intracellular pathways are known to occur (Habib et al. 2020; Escartin et al. 2021). Secondly, the primary norepinephrine nucleus in the brain, locus coeruleus, is known to be affected early in AD, both in humans and in animal models (Weinshenker 2018; Jacobs et al. 2021; Braak and Del Tredici 2011), and perturbed noradrenergic function is likely an important factor both in disease progression in AD patients and accounting for the cognitive decline of AD patients (Weinshenker 2018; Peterson and Li 2018; Holland, Robbins, and Rowe 2021). Restoring noradrenergic signalling in AD mice with perturbed noradrenergic signalling pharmacologically or by pharmacogenetics have been demonstrated to be beneficial (Holland, Robbins, and Rowe 2021). In our study we find a largely retained level of pupil response in the AD mice during startle responses, which under physiological circumstances would suggest that the noradrenergic system was working normally, as a strong correlation between locus coeruleus activity and pupil responses have been established (Reimer et al. 2016; Costa and Rudebeck 2016). The underlying connectivity that enables the pupils to be faithful readouts of the noradrenergic system is to the best of our knowledge not fully established, even though spinal projections from locus coeruleus have been been demonstrated (Hancock and Fougerousse 1976; Costa and Rudebeck 2016; Liu et al. 2017). Our finding of a decoupling between astrocytic Ca^2+^ signalling and arousal-induced pupillary dilation, could be due to perturbed connectivity between locus coeruleus and the relevant nuclei or projections controlling activity in the superior cervical ganglion and consequently the sympathetic projections to the pupils. Lastly, our observations could be explained by a combination of these factors, namely altered intrinsic astrocytic responses in reactive astrogliosis combined with perturbations in the correlation between locus coeruleus and pupillary responses. Future studies are warranted to pinpoint the molecular mechanisms at play.

Previous studies have shown mixed results regarding an effect of plaque-astrocyte distance, with reports of both hyperactivity close to plaques (Delekate et al. 2014) and no such correlation except for plaques serving as initiation sites for pathological intercellular Ca^2+^ waves (Kuchibhotla et al. 2009). In our awake-behaving mice, we found no clear correlation between Ca^2+^ activity and 3D distance reconstruction of plaque positions relative to the imaging plane (Fig. 1E). The apparent discrepancies in the litterature and the present study may be due to different AD mouse models, different age groups investigated, different Ca^2+^ indicators employed or lastly the effects of removing anesthesia allowing for a much richer repertoire of astrocytic Ca^2+^ signalling to emerge, effectively masking a potential weak correlation.

Astrocytes have a highly complex and specialized morphology, and this morphology is known to change in reactive astrogliosis (Escartin et al. 2021). The majority of astrocytic processes are much smaller than what can be clearly delineated by non-super resolution optical microscopy, but to what extent these small processes are altered in reactive astrogliosis is unknown, although the astrocytic territories are known to be preserved (Wilhelmsson et al. 2006). We cannot rule out that gliosis-induced morphological changes in combination with potential subtle differences in GCaMP6f expression and elevated baseline intracellular Ca^2+^ concentration (Kuchibhotla et al. 2009) may influence our results. However, the ability for the whole FOV to be activated during spontaneous running and startle responses and the lack of correlation (if anything a negative correlation) between distance to the nearest amyloid plaque and gross level of Ca^2+^ signaling suggests that our findings are not due to degree or pattern of sensor expression.

The present study underscores the importance of studying astrocytic Ca^2+^ signals in unanesthetized mice, and to carefully consider animal behavior when interpreting astrocytic Ca^2+^ dynamics. By lifting the confounding effects of anesthesia we found that the astrocytic hyperactivity previously reported in Alzheimer’s disease mouse models was only a part of the total picture. At first glance, the physiological Ca^2+^ responses were remarkably well preserved, and not characterized by a general increase in astrocytic Ca^2+^ signalling. However, behavior like quiet wakefulness and locomotion are not static entities, and the degree of activation of all relevant parameters needs to be taken into account with statistical modelling to be able to conclude if there are relevant differences between the genotypes. We were able to demonstrate attenuated Ca^2+^ dynamics and an uncoupling between astrocytic Ca^2+^ signalling and arousal in the tg-ArcSwe mice, and given the growing spectrum of roles ascribed to astrocytic Ca^2+^ signalling in higher brain functions, the present findings may highlight one cause for the cognitive decline of AD patients.

## MATERIALS AND METHODS

### Animals

Tg-ArcSwe mice carry a human AβPP cDNA with the Arctic (p. E693G) and Swedish (p. KM670/671NL) mutations where the human AβPP gene is inherited only from male mice to ensure a more uniform onset of Aβ-deposition (Lillehaug et al. 2014). The transgenic animals develop parenchymal Aβ-plaques from 6 months of age, and cerebral amyloid angiopathy (CAA) from 8 months of age (Yang et al. 2011; Lord et al. 2006). 6 tg-ArcSwe mice and 5 WT littermates (both males and females) were used in this study. Sample sizes were determined based on our previous studies using similar techniques (no power calculations were performed). No randomization or blinding was performed. Genotyping was performed as previously described with primers annealing to the Thy1-promoter and the human APP-transgene (Lord et al 2006). The aminals were housed under standard conditions at a professional veterinarian facility with 12-hour dark-light cycles and unrestricted access to food and water. All animal procedures were in accordance with the National Institutes of Health Guide for the care and use of laboratory animals and approved by the Norwegian Food Safety Authority (project number: FOTS #11983).

### Viral transduction and delivery of fluorophores

Serotype 2/1 recombinant adeno-associated virus (rAAV) from plasmid construct pAAV-GFAP-GCaMP6f (Chen et al. 2013) was generated (rAAV titers about 1.0–6.0 × 10^12^ viral genomes/mL) and used for visualizing astrocytic Ca^2+^ signalling. GCaMP6f was amplified by PCR from pGP-CMV-GCaMP6f with 5′ *Bam*HI and 3′ *Hin*dIII, and sub-cloned into the recombinant rAAV vector pAAV-6P-SEWB (Shevtsova et al. 2005) for generating pAAV-*SYN*-GCaMP6f. The human glial fibrillary acidic protein (*GFAP*) promoter (Hirrlinger et al. 2009) was inserted with *Mlu*I and *Bam*HI into pAAV-*SYN*-GCaMP6f construct for obtaining pAAV-*GFAP*-GCaMP6f. Serotype 2/1 rAAVs from pAAV-*GFAP*-GCaMP6f was produced (Tang et al. 2015), and purified by AVB Sepharose affinity chromatography (Smith, Levy, and Kotin 2009), following titration with real-time PCR (rAAV titers about 1.0–6.0 × 10^12^ viral genomes/mL, TaqMan Assay, Applied Biosystems). To visualize amyloid plaques, 7 mg/g methoxy-X04 (Syverstad Skaaraas et al. 2021) dissolved in 0.1 M phosphate buffer saline (PBS), was injected intraperitoneally 24 hours prior to imaging. We found that one injection provided enough signal to outline amyloid plaques for up to three days.

### Surgical preparation

Mice were anaesthetized with isoflurane (3% for initiation, then 1–1.5% for maintenance) in room air enriched with 20% pure oxygen, and given buprenorphine 0.1 mg/kg s.c. preemptively for analgesia. Bupivacain was administered subcutaneously over the skull, and left for 10 minutes before a boat shaped skin flap was removed. After removing the skin, a 2.5 mm diameter craniotomy was drilled over the somatosensory cortex with center coordinates 3.5 mm lateral and -1.5 mm posterior to Bregma. Virus was injected (70 nL at 35 nL/min at 200 µm depth from the brain’s surface) at three evenly spaced locations positioned to stay clear of large blood vessels, and a glass plug consisting of two coverslips glued together were placed in the craniotomy, slightly pressing the dura to prevent dural overgrowth (Bojarskaite et al. 2020). The surrounding area of the skull was sealed with cyanoacrylate glue and a layer of dental cement. Post-operatively the mice were given meloxicam 2 mg/kg for two days. Only animals with normal post-operative recovery were included in the study. Mice were left to recover for a minimum of two weeks before habituation to head-fixation and imaging.

### Two-photon microscopy

After recovery, the animals were imaged in layers 1–3 of the barrel cortex using a two-photon microscope (Ultima IV from Bruker/Prairie Technologies, Nikon 16X, 0.8 NA water-immersion objective model CFI75 LWD 16XW, Spectra-Physics InSight DeepSee laser, Peltier cooled photomultiplier tubes model 7422PA-40 by Hamamatsu Photonics K.K.). An excitation wavelength of 990 nm was used for GCaMP6f imaging, and separate recordings were made using 790 nm excitation light for imaging methoxy-X04, and 890 nm for Texas-Red labelled dextran for visualization of the vasculature. Image time series were recorded at 30Hz. 3D volume recordings of the morphology and plaque locations were performed in a subset of experiments. All images were recorded at 512×512 pixels with a resolution of approximately either 0.42 or 0.67 μm per pixel (FOV of 215×215 μm, or 343×343 μm, respectively). For imaging, the mice were head-fixed to a custom build stage that allowed free locomotion on a wheel attached to a rotary encoder. During experiments the mice were monitored with an IR sensitive camera, illuminated by an IR LED diode. A second camera was used to capture the pupil dynamics (see below). Air-puffs to elicit startle responses were delivered by a Picospritzer III (Parker). Instrument synchronization and data acquisition were performed with a custom-made LabVIEW 2015 (National Instruments) virtual instrument.

### Behavioural analysis

Mouse locomotion was recorded with a rotary encoder connected to the running wheel. Locomotion signal was captured using a National Instruments data acquisition card using a counter task in the NI Max software, activated through a custom LabView (2015) VI. Data was processed with custom MATLAB scripts to classify running and quiet wakefulness. Criteria for run and quiet wakefulness episodes were validated by manual observation, and defined as follows: Running was defined as continuous segment of at least 4 s forward wheel motion at over 30 degrees/s, no movement faster than 20 degrees/s the last 10 seconds prior to start of running. Spontaneous running was not defined within 30 seconds following air-puff. Quiet wakefulness was defined as continuous segment of at least 10 s duration with less than 2 degrees/second locomotion, as well as no locomotor activity faster than 2 degrees/s for 15 seconds before segment start. Quiet wakefulness episodes were not defined within 30 seconds following air-puff. Quiet wakefulness periods were reviewed manually through the IR-surveillance video recordings to ensure animals were awake when sitting still.

### Pupillometry

Pupil size was recorded with a Basler Dart USB camera (daA1600-60um) with a 25mm fixed focal length lens and 2X fixed focal length lens extender (Edmund optics, items #59-871 and #54-356). The pupil was back-illuminated with the spillover light from the two-photon microscope laser coming through the cranial window (Yüzgeç et al. 2018). As two-photon imaging must be conducted in the dark where the pupil would be fully dilated, we illuminated the mouse eye contralateral to the recording side using 470 nm blue light fiber, with a shielding to avoid light contamination, to slightly constrict the pupil. Pupil size was manually delineated for time periods of isolated runs and startle using a custom build tool developed in MATLAB 2020a. The ratio of pupil to eye-size ratio was calculated and used in the analysis.

### Image processing and analysis

Recordings were exclusively processed using MATLAB 2018a to 2020b. We used our recently published imaging analysis toolbox ‘Begonia’ to remove motion artefacts, mark ROIs and perform event-based Ca^2+^ signal detection (regions-of-activity method; ROA). Methods and algorithms related to this processing pipeline are described in detail in Bjørnstad et al. 2021 (Bjørnstad et al. 2021). Astrocytic Ca^2+^ signals were detected using the ROA method, and the ROA density (i.e. fraction of the compartment analyzed being active) and ROA frequency in the whole FOV or in anatomical subcompartments were calculated.

### 3D amyloid plaque mapping

Before any recording, z-stacks of images at 5 μm intervals taken at 512 × 512 pixels were recorded while illuminating the tissue with 790 nm laser light. The stacks started approximately at or below dura mater and extended 100–200 μm down. All other recordings were undertaken inside this mapped volume to ensure plaques outside the imaging plane were accounted for. For each imaged time series inside the volume we recorded an additional single image at 790 nm to ensure the precise location of plaques were known. Plaque locations were detected by binarizing the stack at a manually set threshold at which the morphology of plaques were visible. The binarized single images were used to manually align the 2D time series data with the 3D plaque volume. The resulting 3D binary image had small points removed using the Matlab function bwareaopen.

### Tissue processing

All animals were sacrificed after final imaging procedures, at the age of 18 months. The animals were anesthetized with Isofluran Baxter (IsoFlo, Abbot Laboratories) and intraperitoneally injected with ZRF mixture (Zolazepam 3.8 mg/ml, Tiletamin 3.8 mg/ml Xylazine 0.45 mg/ml and Fentanyl 2.6 µg/ml) before transcardial perfusion with 4?C 2% dextran in 0.1 M phosphate buffer (PB) for approximately 30 seconds, immediately followed by 4% formaldehyde (FA) in PB for 10 minutes at the speed of 6 ml/min. Following perfusion, the brains were extracted and post fixed by immersion in the fixative at 4?C overnight protected from light. The tissue was stored in 0.1 % FA in PB at 4?C protected from light until further processing. Cryoprotective steps in graded sucrose solution (10%, 20%, and 30% sucrose in PB) were performed before the brains were cut on a freeze microtome (Thermo ScientificTM Microm KS 34) in 40 µm free floating coronary sections and stored in 0.1 % FA in PB at 4?C protected from light until usage. In addition, 8 tg-ArcSwe animals and 8 wild type littermates were sacrificed at 12 months of age and used for qPCR analysis. These animals were anesthetized as described above and decapitated. The brains were extracted, and the left hemisphere was dissected into the frontal cortex, hippocampus, cerebellum and the rest of the brain. This tissue was frozen and stored in -80?C pending analysis.

### RNA isolation and real time PCR

48 hours prior to RNA extraction, the samples were suspended in RNAlater(tm)-ICE (Ambion; Cat#: AM7030). To isolate total RNA from the frontal cortex tissue samples, the RNeasy Mini Kit (QIAGEN, Hilden, Germany), including the on column DNase digestion, was used. The RNA concentration and integrity were determined using a NanoDrop 2000c spectrophotometer (ThermoFisher Scientific) and ethidium bromide visualization after agarose gel electrophoresis, respectively. Following the manufacturer’s protocol, 1 µg of total RNA was reverse - transcribed into cDNA with Oligo (dT)15 using the GoScript Reverse Transcription System (Promega, Madison, USA, Cat#: A5001). All the cDNA samples were diluted in Tris-EDTA buffer (pH 8.0) to a final concentration of 2.5 ng/μl. Real-Time PCR was carried out in a total volume of 20 µl, containing 2x AB Power SYBR® Green PCR Master Mix (ThermoFisher Scientific) with gene specific primers (at a final concentration of 200 nM) and 2 µl cDNA samples. Amplification was performed on the StepOnePlus system (Applied Biosystems) with the following conditions: 95°C for 10 minutes, followed by 40 cycles of 95°C for 15 seconds and 60°C for 1 minute, followed by melting curve analysis to check for unspecific products. Each sample was run in duplicates. Using the NormFinder software (Andersen et. al. 2004), *HPRT1* was determined as an internal control for normalization of the gene expression. The primers were designed online using Primer-BLAST and setting the amplicon size to a maximum of 200 bp. The primers designed span exon-exon junction, and standards prepared as previously described (Rao et al. 2021). Details of the *Gfap* forward and reverse primer are presented in Table 1.

**Table 1.**
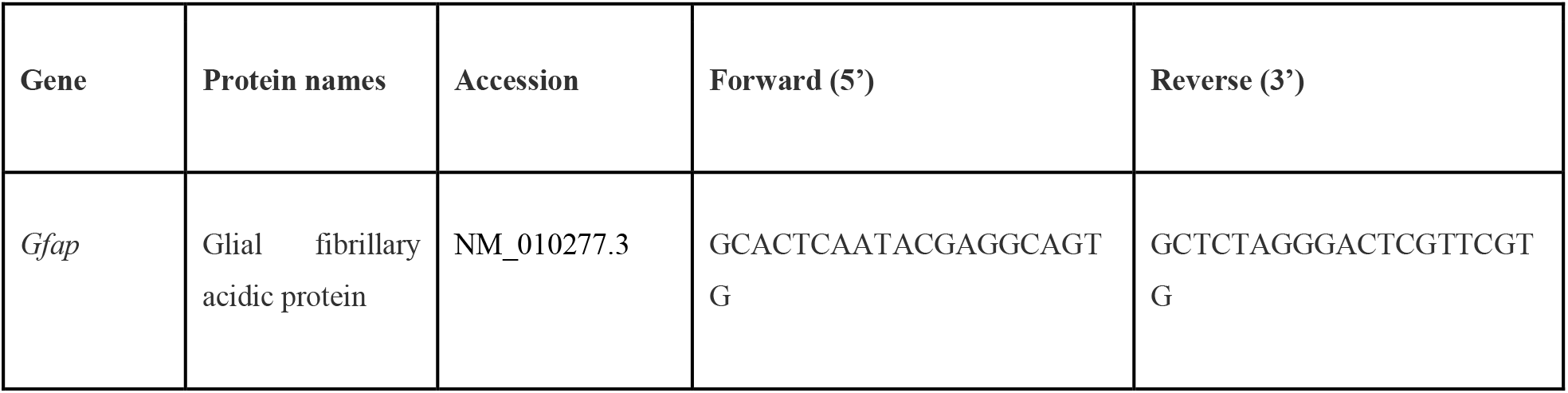
Primer used for mRNA quantification.

### Immunohistochemistry

One section from each animal was chosen for quantification of astrogliosis and washed in PBS 0.01 M for 10 minutes, followed by two times in 0.1% TritonX100 in PBS (PBST) for 5 minutes. The PBST was removed and blocking (10% normal donkey serum (NDS), 1 % bovine serum albumin (BSA), 0.5% Triton X100 in PBS) performed for one hour at room temperature. This was directly followed by incubation overnight at room temperature with primary antibodies (GFAP; host: mouse; diluted 1:1000; Sigma-Aldrich; Cat# MAB360. GFP; host: chicken; diluted 1:2000; Abcam; cat# ab13970) diluted in antibody solution (ABS; 3% NDS, 1 % BSA, 0.1 % Triton X100 in PBS). The following day the sections were rinsed in 0.1% PBST two times for 1 minute, followed by three times for 10 minutes. Secondary antibodies (CY5 Donkey Anti-Mouse; Jackson ImmunoResearch Labs; Cat# 715-175-151. CY3 Donkey Anti-Chicken; Jackson ImmunoResearch Labs; Cat#703-165-155) were spun in a centrifuge for 10 minutes at 13000 rpm, diluted 1:500 in ABS, and the sections incubated for 1 hour at room temperature. After the second incubation, the sections were washed in PBS for 10 minutes, three times. Propidium iodide (diluted 1:5000 in 0.01 M PBS; Sigma Aldrich; Cat # 04511 (Cellstain double staining kit)) for nuclear staining was added for 10 minutes, before rinsing the sections twice for 5 minutes in PBS. All sections were transferred to distilled water and mounted with ProLong Gold antifade reagent (ThermoFisher Scientific; Cat# P36934). They were stored in -20 ?C protected from light until confocal imaging. For electron microscopy: EM standard procedure was followed for embedding and preparing of the tissue (Yang et al. 2011). For enhancing the contrast, uranyl acetate (Fluorochem) in double distilled water and lead citrate was used. The sections were examined in a transmission electron microscope (TECNAI 12).

### Confocal imaging

GFP-positive astrocytes (indicating GCaMP6f expression) were used to locate the appropriate cortical area used for in vivo imaging. Only sections where we could locate positive GFP staining within the cortex corresponding to the image area were chosen for confocal imaging and further analysis. All single plain and z-stack images were acquired using a Zeiss LSM 710 confocal microscope. To provide overview and verify that the correct area was identified, one tile scan of 3072 × 3072 pixels was achieved with a 40x objective (1.20; water korr M27) using 3 channels (CY2, CY3 and CY5) for wild types and 4 channels (Dapi, CY2, CY3 and CY5) for tg-ArcSwe. Next, one z-stack of 2048 × 2048 pixels (40x objective; 1.20; water korr M27) from within the GFP positive area was acquired from cortical layer 2. All single plane and z-stack images were obtained with identical settings. In addition, a z-stack of 1024 × 1024 pixels was obtained (40x objective; 1.20; water korr M27) outside the located GFP positive stained area. No post-processing on the analysed images were performed.

### 3D reconstruction analysis of astrocytes

This procedure was adapted from the details outlined in Tavares et al. (Tavares et al. 2017) that utilize the free software plugin Simple Neurite Tracer (SNT) of FIJI-ImageJ (Longair, Baker, and Armstrong 2011). Only astrocytes with a single nucleus where at least ⅔ of the circumference was covered by GFAP staining were selected for 3D reconstruction. Astrocytes with processes touching the borders of field of view were omitted. 8 randomly selected astrocytes from each z-stack (2 from each image quadrant) obtained within the GFP positive imaging area were used for analysis. To quantify and visualize the morphological complexity of the astrocytes, we analysed the number of processes, total length of processes in µm, process thickness (µm^3^) and number of intersections (provided from the Sholl analysis).

### GFAP area fraction analyses

Z-projections based on the CY5-channel to visualize GFAP positive astrocytes of all stack images were rendered using the FIJI-ImageJ software (Schindelin et al. 2012). The images were blinded to the analyst, converted to 8 bit images and the scale removed. The threshold was manually adjusted so that only what was considered to be GFAP positive staining was red, and a percentage value of positive staining in the image was obtained.

### Statistical analysis

Astrocytic Ca^2+^ signals were studied by means of the ROA density – a number between 0 and 1 indicating the fraction of the compartment with activity. Here, the area may be the entire field-of-view (FOV), or we may limit ourselves to the area identified as belonging to cellular subcompartments; the astrocytic processes, somata or endfeet. In each trial we had time series of ROA density lasting around 300 s (approximately 9000 frames). Within each trial, the startle period was defined as starting from the air puff at 150 s and lasting 600 frames (∼20 seconds). In addition, one or more time periods within the trial could be identified as spontaneous runs, or as periods of quiet wakefulness. A first important question concerns the choice of summary statistics adequately describing the Ca^2+^ response in such periods of interest (runs and startle), which are dynamic behavioral states that entail both acceleration, steady locomotion and deceleration. We have studied the mean and max ROA density and the ROA density rise rate which is defined as the *maximal increase in ROA density over a maximum of 50 frames*. Initial explorations indicated that the main results are fairly robust to the choice of window length, and 50 frames appeared to be a sensible choice compared to the kinetics of astrocytic Ca^2+^ signals. The ROA density rise rate is meant to capture some of the dynamics in astrocytic Ca^2+^ signaling, and can be understood as the maximum acceleration inside the time period of interest. The rise rate has a high correlation with the maximum, and if the length of the window is increased sufficiently these two statistics tend to become almost identical (since most traces are close to zero at some point in the trial). See the Supplementary Fig. 5A which displays both the max ROA density and the rise rate in an example.

Pupil size measurements had a coarser time resolution than for the astrocytic Ca^2+^ signals; see example in Supplementary Fig. 5B, showing the pupil sizes around a startle response. The pattern in the figure – a sharp increase in pupil size after air puff, before a gradual decline – was quite typical. Therefore, we chose to only consider the measurements in a time-window of 6.67 seconds on each side of the airpuff (or start of running). We defined the *pupil dilation* as the relative increase in the ratio of pupil diameter to eye diameter, inside this window. In other words, we calculate the average ratio before the air puff (or start of running) and after the airpuff, compute the difference and divide by the average before the air puff.

#### Interpreting hierarchical plots

We have chosen to present some of our data in the form of *hierarchical plots* (see Figure 2B and C, Figure 3B and C, and Supplementary Fig. 1, 2 and 3). These plots allow the reader to assess the degree of separation between the genotypes and at the same time get an impression of the *variation at different levels of the analysis* – in our case, the variation between different mice of the same genotype and the variation between repeated measurements on the same mouse. At the lowest level – the trial level – we have points representing the observations themselves. For the spontaneous runs, there are sometimes more than one run per trial and in that case the points are the median ROA density rise rate in these runs. The maximal number of spontaneous runs in a single trial was 5 (average: 1.6 runs per trial). At the middle level we have the median ROA density rise rate for each mouse, and at the top level the median ROA density rise rate for each genotype. The lines between the levels indicate which observations belong to each mouse, and to each genotype respectively. We have made similar plots for the max ROA density also. The hierarchical plots are a useful tool for exploratory analysis. They are also meant to promote transparency in scientific reporting, and to highlight the importance of intra-group variation. Still, it is important to realize that the impression conveyed by the plots might not be identical to the results from statistical modelling. In the plots, we do not include the influence of various technical and biological covariates which one typically would include in a statistical model. Some of the variation between observations belonging to the same mouse and between different mice of the same genotype may be explained by such covariates, as we will see in the next section.

#### Modelling

Statistical analyses were conducted in R (version 4.1.1). The ROA density rise rate was modelled by linear mixed effect regression models which were fitted using the glmmTMB package (Brooks, Kristensen, and Van Benthem 2017). We conducted two sets of analyses: (i) to investigate potential differences in ROA density rise rate between the two genotypes; (ii) to investigate the relationship between ROA density rise rate and the pupil dilation, including potential differences between the two genotypes with respect to this relationship. For (i) the coefficient of primary interest is the one belonging to the genotype variable, while for (ii) we are interested in the effect of pupil dilation on the ROA density rise rate, as well as the interaction between this pupil effect and the genotype. In both of these sets of analyses we adjusted for the following fixed effect covariates: the level of optical zoom (2 levels), the depth of the measurements (in μm) and the maximal speed in the relevant time window. We included random intercepts for each mouse (5 WT and 6 tg-ArcSwe). We analyzed the ROA density max and mean values with similar models as the ones described here for the rise rate. For (i), the number of observations ranged between 77 to 117 trials with startle data (depending on the subcompartment) and between 72 and 109 episodes of spontaneous running, while for (ii) the number of observations ranged between 60 and 86 for the startle data and between 35 and 44 for the spontaneous runs. There were less observations for the (ii) analyses because some episodes/trials had missing or incomplete pupil measurements.

The sensitivity of our result to these modelling choices were assessed by various robustness checks, see next section. The adequacy of model assumptions was investigated by residual plots (Hartig, n.d.). In the cases where the residual plots indicated deviations from the assumption of constant residual variance, we extended the model by allowing the residual variance to vary as a function of genotype. The reported p-values are based on the t-distribution, with degrees of freedom as provided from the glmmTMB package. No corrections for multiple comparisons were applied.

When analysing the relationship between distance from the nearest plaque and astrocytic Ca^2+^ signalling (Figure 1E) we considered the mean ROA density in each ROI (n=10988) in the quiet wakefulness episodes. If present, any effect of plaque proximity on Ca^2+^ signalling should be discernible among ROIs observed in the same episode. The dashed lines in Figure 1E show the effect of distance on Ca^2+^ signalling within each episode, and they form an uncertain picture: in some episodes there is a weak positive relationship, with seemingly higher mean ROA density further away from plaques, while in many episodes there is a negative relationship, with somewhat higher mean ROA density close to plaques. The overall line is found by fitting a linear mixed effect model with mean ROA density as the response, with a fixed effect of distance and with each episode having its own random intercept and slope for the distance effect.

For the results presented in Figure 4: Unless otherwise stated, the data are presented as mean ± standard error of the mean (SEM). A p-value equal to or below 0.05 was considered statistically significant. Mann-Whitney U-test was used to analyse the number of processes, total length of processes in µm, process thickness in µm^3^ and area fraction of positive GFAP staining. Two-way ANOVA followed by Sidak post hoc comparison was used to analyse the number of intersections. Statistical analysis was performed in GraphPad Prism version 8.0.1 for Windows (GraphPad Software). For qPCR analysis, mean copy number per ng of total RNA was compared between genotypes by Mann-Whitney U test in SPSS Statistics 26 (SPSS).

#### Robustness checks

In order to check the robustness of the various statistical analyses, the stability of estimates and p-values was examined with respect to the length of window, for the rise rate response variable, and also sensitivity to individual mice and trials.

The type of sensitivity analysis performed is illustrated (see Supplementary Figure 6) for the analysis of the uncoupling between pupil dilation and astrocytic Ca^2+^ responses (see Figure 3 for details about the data). Again, for the purpose of illustration, we focus on the interaction between pupil dilation and genotype for the ROA density rise rate response variable; see Supplementary Figure 6 for details. Similar analyses were performed for the main statistical models, none of these sensitivity analyses provided clear or strong evidence for any change in the main conclusions.

## Data and source code availability

A complete dataset of raw and processed data including Ca^2+^ signal traces, behavioral monitoring traces (all sampled and time aligned to 30Hz) and pupil tracking data are provided at: https://doi.org/10.11582/2021.00100. Data analyses were performed with the Begonia toolkit (Bjørnstad et al. 2021), which is available at https://github.com/GliaLab/Begonia.

## Acknowledgements

This work was supported by the Olav Thon Foundation, the Letten Foundation, Norwegian Health Association, Alzheimer fond, the Research Council of Norway (grants #249988), the South-Eastern Norway Regional Health Authority (grant #2016070). We acknowledge the support by UNINETT Sigma2 AS for making data storage available through NIRD, project NS9021K. Prof. Lars Lannfelt is gratefully acknowledged for help to the development of the tg-ArcSwe mouse model at Uppsala University.

## Author contributions

Conceptualization: E.A.N., R.T., R.E.; Methodology: K.S.Å., G.H.E.S.S., D.M.B.,R.T., R.E., V.J., W.T., L.N.G.N; S.B.R. Software: K.S.Å., D.M.B., R.E., C.C., G.H.H.; Formal analysis: K.S.Å., G.H.E.S.S., R.E., C.C., G.H.H.; Investigation: L.B.; Resources: W.T., E.A.N., R.T., R.E.; Writing— original draft: O.O., R.E., K.S.Å., G.H.E.S.S.; Writing—review and editing: O.O., K.S.Å., G.H.E.S.S., R.T., D.M.B., C.C., G.H.H., K.M.G.B., V.J., L.N.G.N; Visualization: K.S.Å., G.H.E.S.S., R.E.; Supervision: R.E., E.A.N., V.J., R.T.; Funding acquisition: E.A.N., R.E., R.T.

## Supplementary Information

**Supplementary Figure 1.**
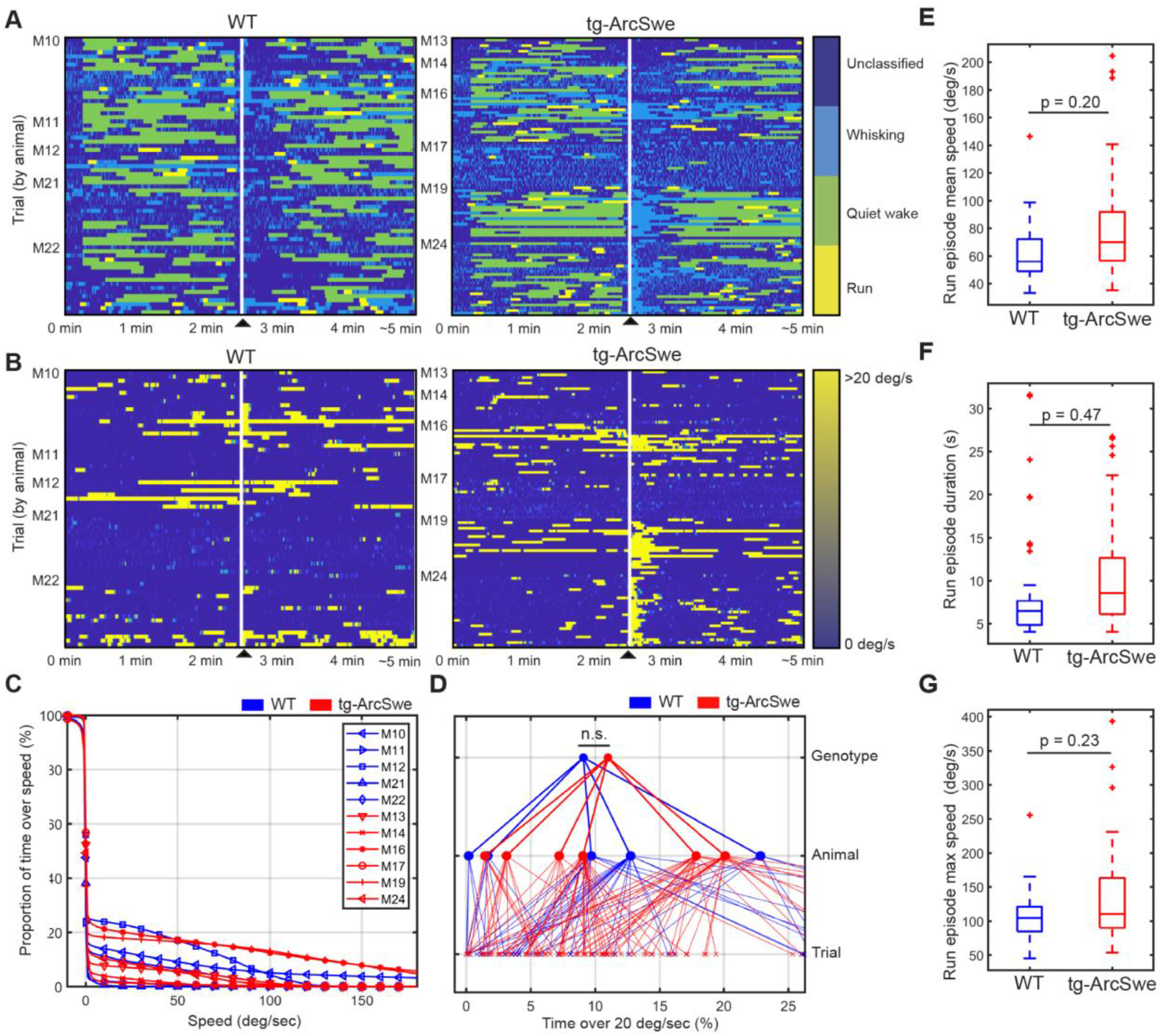
Locomotor behavior in WT and tg-ArcSwe mice. ▲ indicates air-puff. (A) Behavioral classification for each individual trial, grouped by mouse. See methods for classification and filtering algorithm used. (B) Running wheel speed for each individual trial, grouped by mouse. 20 degrees of rotation of the running wheel per s was the threshold used to classify running. (C) Percentage of total time a mouse spent above a given speed across all trials. (D) Percentage of total time a mouse was running (> 20 degrees/s). Raw running data not filtered by the run classification algorithm (p = 0.35). (E, F, G) Post-classification run episode characteristics.

**Supplementary Figure 2.**
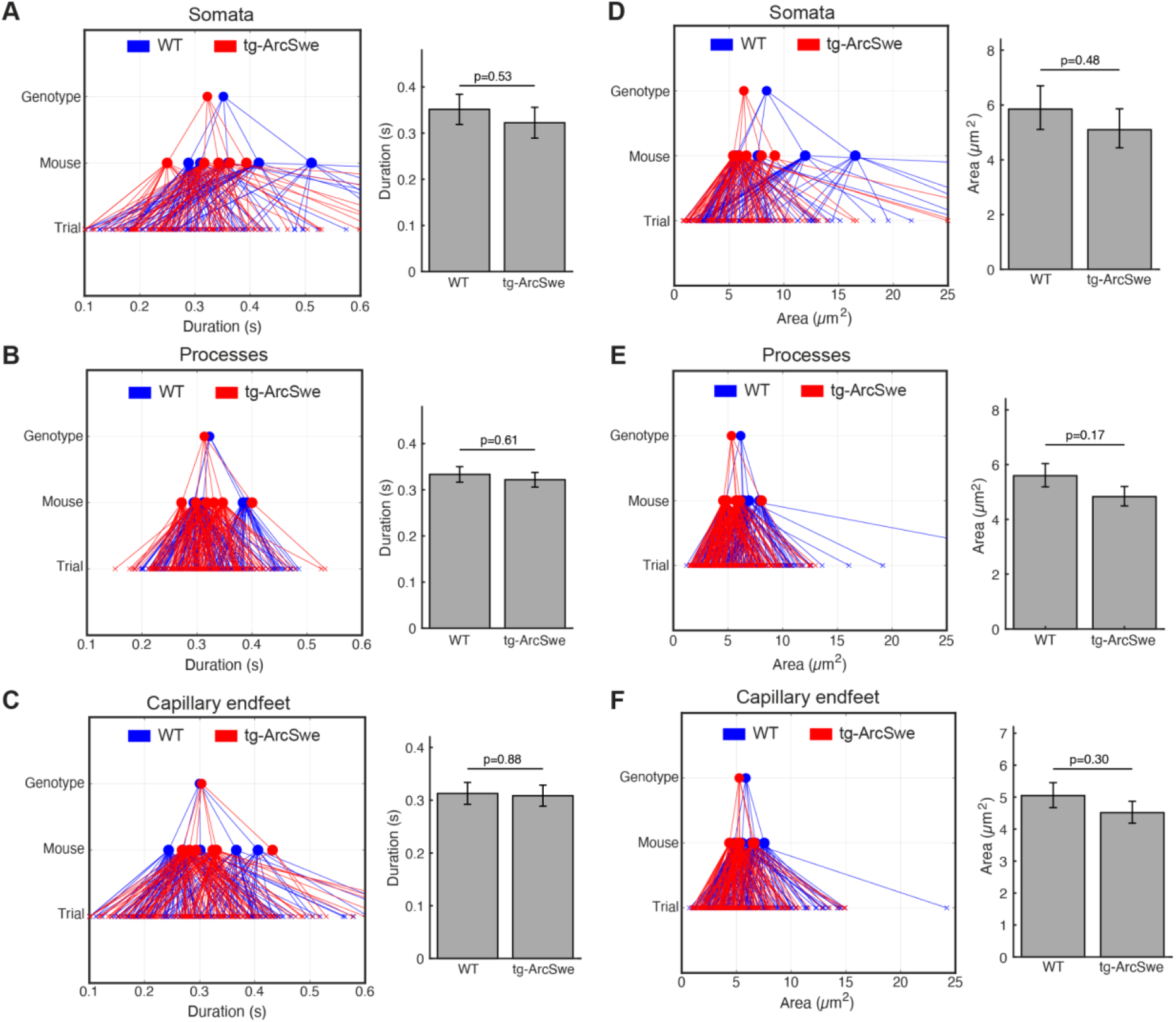
Regions-of-Activity during quiet wakefulness exhibit the same characteristics in both genotypes. During quiet wakefulness episodes, active regions detected by the ROA algorithm. Compartment association is determined by the centre of the region-of-activity. (A, B, C) Mean duration of active regions during quiet wakefulness in different cellular compartments. Right: GLME statistical analysis estimates. (D, E, F) Mean maximum size of a detected region. Right: GLME statistical analysis estimates.

**Supplementary Figure 3.**
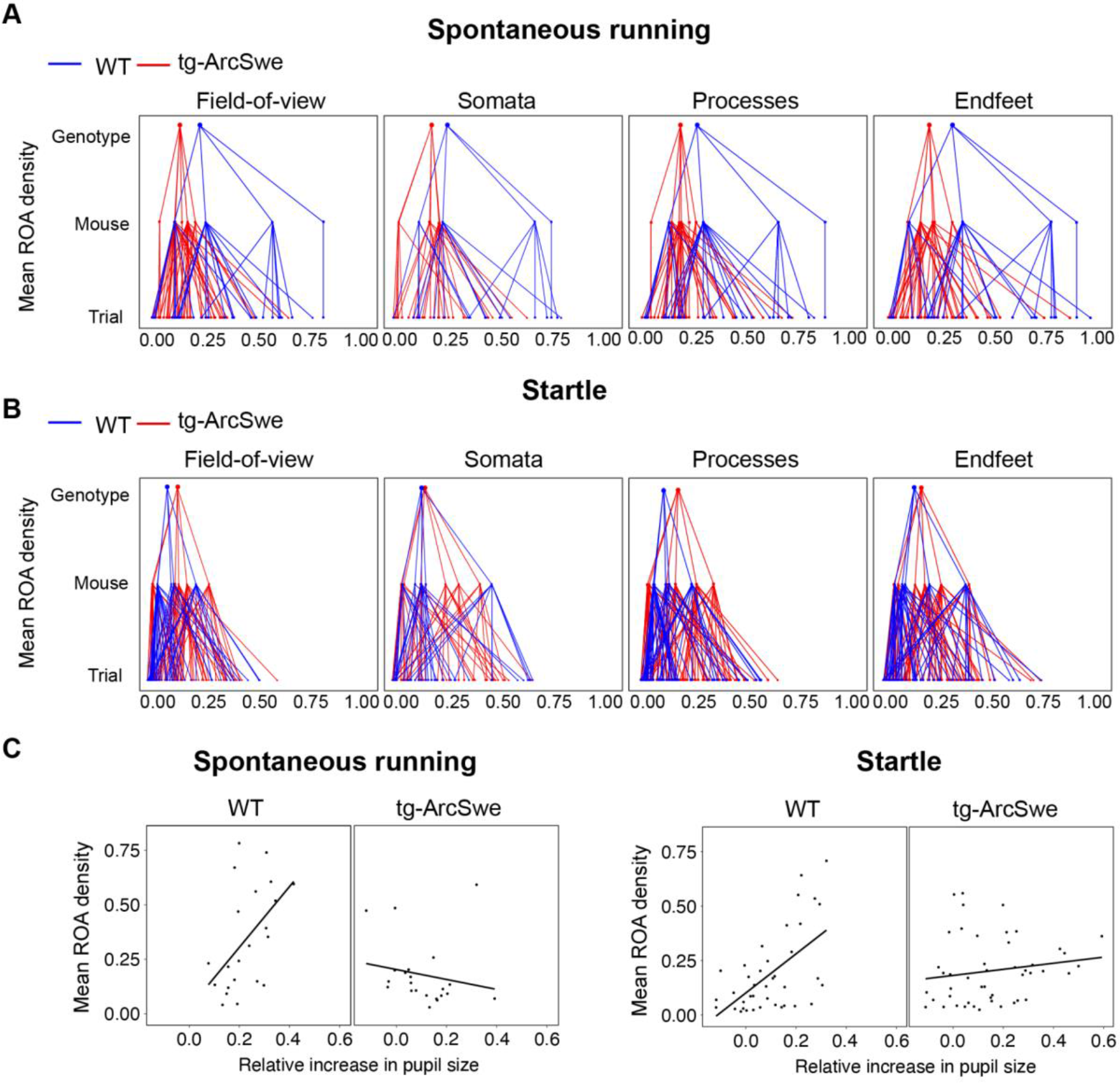
(A) Hierarchical showing medians of mean ROA density per trial, mouse and genotype in the FOV and astrocytic subcellular compartments during spontaneous runs. (B) Same as (A) but during startle responses. (C) Scatterplots of mean ROA density vs. relative increase in pupil size in spontaneous running and startle responses.

**Supplementary Figure 4.**
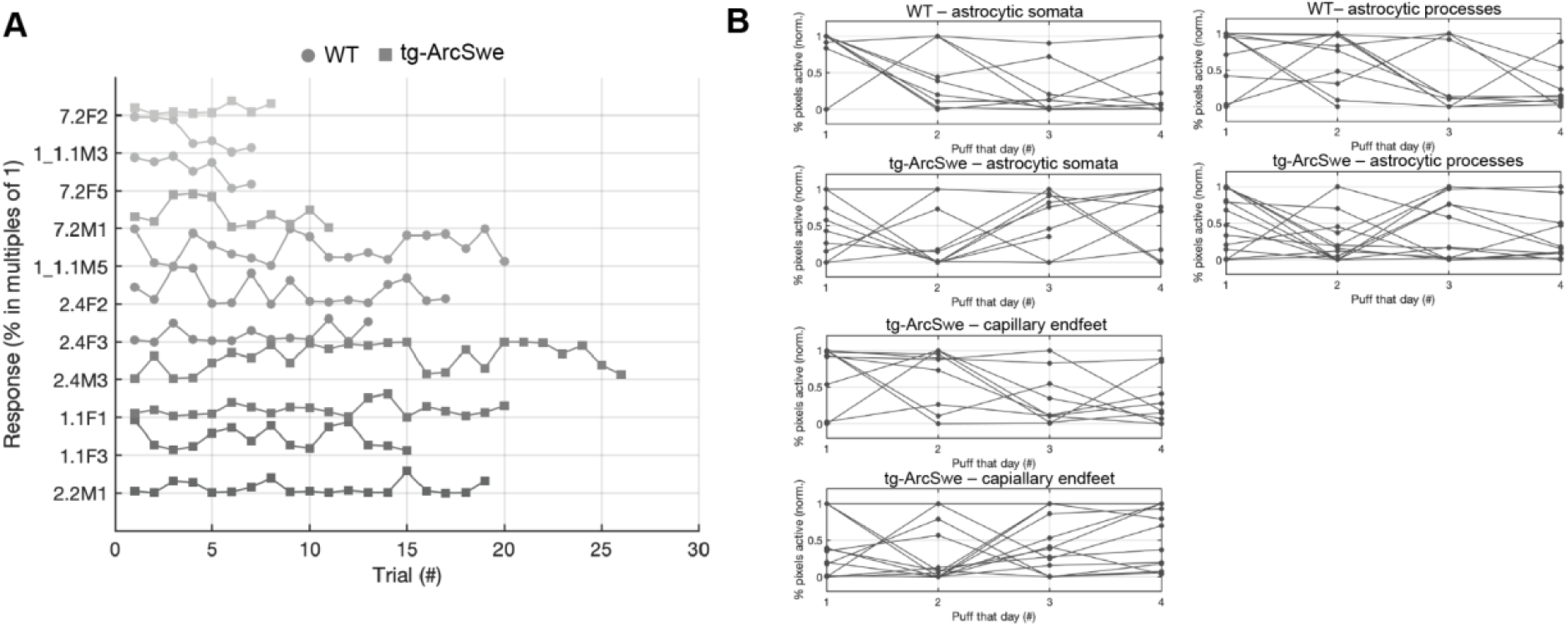
(A) Neither tg-ArcSwe or WT mice displayed any noticeable habituation to air puffs across all trials and experimental days. (B) Within a single experimental day, there was no habituation to air puffs across the different trials.

**Supplementary Figure 5.**
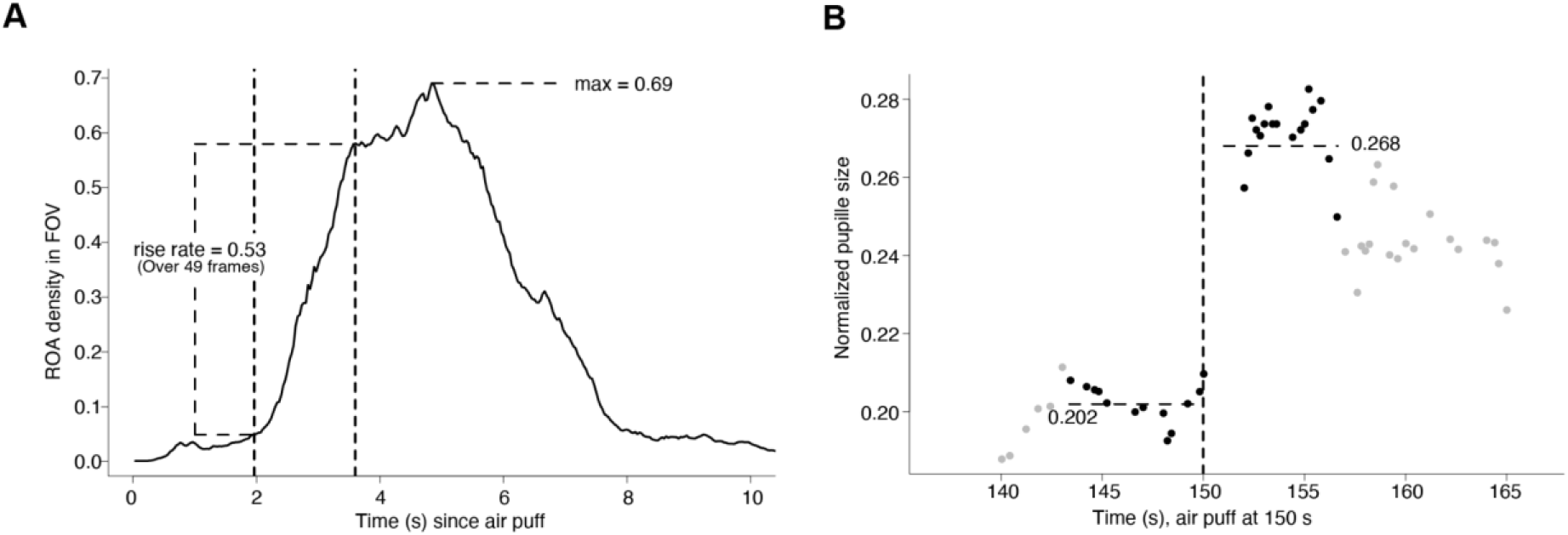
(A) Computation of the ROA density rise rate, and max ROA density, in particular startle period. The vertical dashed lines indicate the time window which had the maximal increase in ROA density, which in this example took place over 49 frames. (B) Computation of the pupil dilation in a particular startle period, here we find (0.268-0.202)/0.202=0.33.

**Supplementary Figure 6.**
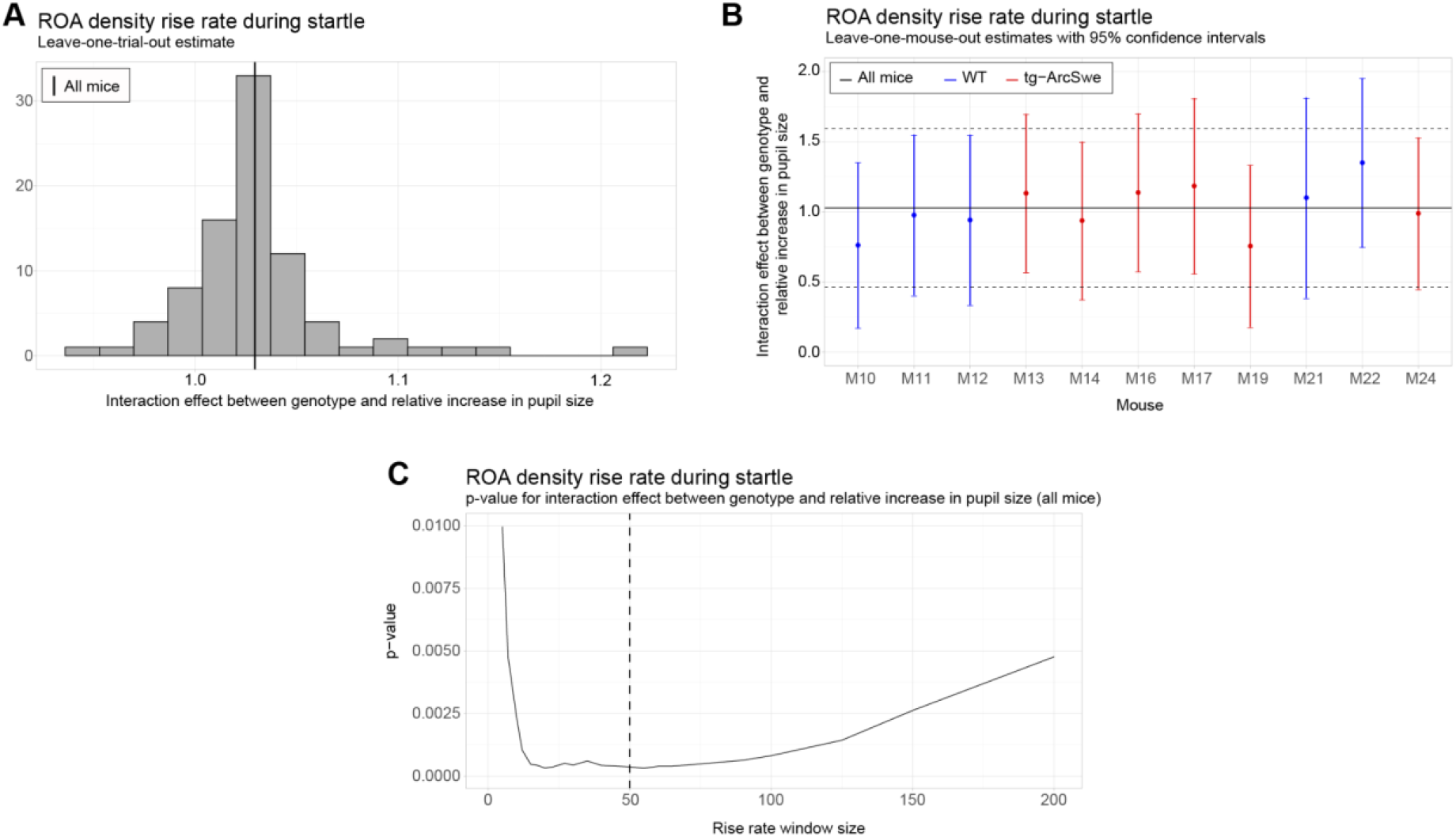
Robustness and sensitivity analyses for the uncoupling between pupil dilation and the ROA density rise rate. All analyses are with respect to the interaction effect between pupil dilation and genotype. (A) Leave-one-trial-out analysis. The full model is repeatedly estimated removing one trial at the time from the dataset (as a type of jackknife resampling). The estimated interaction coefficients are shown as a histogram. Note that the histogram indicates less variation in the estimated interaction when comparing the variance or empirical quantiles of the histogram with the corresponding confidence interval obtained from the linear mixed effect model (see solid and dashed lines in Plot B). (B) Leave-one-mouse-out analysis. In each iteration, one mouse, M10-M24, is removed from the estimation of the model. The estimated interaction effects with corresponding confidence intervals are then compared to the estimate and confidence interval from the full model (solid and dashed horizontal lines). Some mice influence the results more than others, but there appears to be no systematic bias associated with a single mouse. (C) The p-value for the interaction effect for different window lengths. The window length is measured in number of frames. It is clear that the significance of the interaction effect does not depend strongly on a particular choice of window length. In the analysis we used a window of length 50 frames.

